# The Fc fragment of IgMs binds C1q to activate the first step of the classical complement pathway, while inhibiting complement-dependent cytotoxicity

**DOI:** 10.1101/2024.07.10.602503

**Authors:** Andrea J. Pinto, Anne Chouquet, Isabelle Bally, Véronique Rossi, Nicole M. Thielens, Chantal Dumestre-Pérard, Renate Kunert, Christine Gaboriaud, Wai Li Ling, Jean-Baptiste Reiser

## Abstract

Soluble type-M immunoglobulins (IgMs), among the most potent activators of the classical pathway, are key mediators of complement-dependent cytotoxicity, which render them promising drug candidates for the development of alternative drugs in treating autoimmune or inflammatory diseases. In this study, we investigated the biochemical and *in vitro* functional properties of recombinant fragments from IgMs corresponding to the Fc-core in their pentameric or hexameric forms. Biophysical experiments confirmed the crucial role of the IgM Joining chain (J) in favoring homogenous pentamers, while its absence led to heterogeneous population with a mixture of oligomeric forms. By combining size-exclusion chromatography with mass photometry, isolation of enriched samples with IgM hexamers or IgM pentamers without the J chain was possible. Biolayer interferometry demonstrated that both IgM-Fc forms bind C1q and ELISA showed that they induce the *in vitro* C4b deposition when in solid phase. Additionally, our data confirmed the higher efficacy of IgM hexamers compared to pentamers in activating the first component of the classical pathway. Finally, hemolytic assays demonstrate the ability of IgM-Fc constructs to inhibit Ig-induced complement-dependent cytotoxicity, which is likely made possible by the absence of Fab. These findings suggest a possible mechanism of C1 sequestration in plasma by IgM cores and consumption of the initial complement component C4. Our data thus provide important information for the development of IgM-based anti-inflammatory molecules that target specifically complement activation.

## Introduction

Soluble type-M immunoglobulins (IgM) stand out among Immunoglobulin (Ig) classes with their unique structure and their highest level of oligomeric states. They are primarily secreted by naive B cells, but also in some cases, after B cell maturation, to play significant roles in immune responses to infections such as antibody-dependent cellular cytotoxicity (ADCC), antibody-dependent cellular phagocytosis (ADCP) and complement-dependent cytotoxicity (CDC). Like other immunoglobulin classes, they are composed of a heavy chain (H) and a light chain (L) and are submitted to the well-known VDJ gene rearrangements and the somatic hyper-mutation processes during maturation, which are crucial for antigen specificity and assembly of the variable domains composing their antigen-recognition Fab domains. IgMs structurally differ from the other antibodies in the number of H chain constant domains (comprising 4 domains, Cμ1 to Cμ4) and in their overall assemblies. They are found in sera with a high degree of oligomerization, forming either pentamers ((H_2_L_2_)_5_J), which include the additional Joining (J) polypeptide or hexamers ((H_2_L_2_)_6_), devoid of the J chain (for review (1–5)).

Those unique assemblies are made possible by the formation of the so-called fragment crystallizable (Fc) region or Fc-core, comprising the Cμ2, Cμ3, Cμ4 domains, and the C-terminal tailpiece, connected through an extensive disulfide-bond network. The Fc-core is essential for IgMs to achieve their ADCC, ADCP or CDC effector functions by binding to cellular or soluble immune receptors. Among them, binding to the polymeric immunoglobulin receptor (pIgR) and its secretory component (SC) mediates IgM transport across epithelial layer towards mucosal lumen, aiding in immune tolerance towards commensal bacteria (6). The IgM-type immune complexes (ICs) also regulate the immune system though interactions with two Ig receptors, Fcα/μR and FcμR. Fcα/μR is involved in phagocytosis of ICs by B cells to facilitate antigen processing and presentation to helper T cells in the induction of immune response against T-dependent antigens (7) or to negatively regulate immune responses against T-independent antigens (8). IgMs can also be a carrier of the apoptosis inhibitor of macrophage (AIM/CD5L) to regulate thymocytes apoptosis (9). The AIM-IgM complexes affect the interaction with Fcα/μR expressed on follicular dendritic cells (FDCs), slowing ICs internalization to facilitate antigen presentation to B cells and stimulate antibody responses (10). While Fcα/μR can bind both IgA and IgM subtypes, FcμR (also named FCMR, Toso/Fas apoptotic inhibitory molecule 3 or FAIM3) is only expressed by lymphocytes, specific for IgM, and presumably involved in immune cellular activation and control of autoantibody production (11). Finally, CDC is triggered by IgMs when bound to surface-exposed antigens and recognized by the first component of the classical pathway of the complement system, the C1 complex. They are known to be the most potent activators of the amplifying proteolytic cascade that leads to the production of the main C3 and C5 complement convertases, which regulate the immune system stimulation, and lead to the formation of the membrane attack complex and the elimination of the pathogens or the infected cells (12, 13).

In terms of antigen recognition functions, the IgMs high oligomerization gives them a longer half-life in sera and a higher multivalency than IgGs: 10 to 12 antigen-binding fragments (Fab) are found in IgMs *vs.* 2 in IgGs, leading to a greater avidity of IgMs to neutralize their specific targets. Most of all, pentameric and hexameric soluble IgMs are known to be more potent activators of the CDC and the classical complement pathway than hexameric structures of bivalent IgGs (12, 13).

The IgM specific properties thus make them attractive candidates for the design of new biotechnological tools and future therapeutics. They have already inspired IgM-based or Fc-fusion drug research, although no clinical trials have yet reached completion (3). Among many, examples are IgM-like inhalable ACE2 fusion protein (14) and IgG-IgM Fc chimeras such as the Fc-μTP-L309C (15). However, biotechnical bottlenecks in recombinant production and characterization of IgMs or chimeras, and the functional uncertainties of newly design constructs urge for a deeper understanding of their modes of action and potencies.

In the present study, we are reporting biochemical and functional *in vitro* characterization of fragments from IgMs corresponding to the rigid Fc-core in their two forms, pentameric and hexameric. Using data from biophysical methods, and in-house enzyme-linked immunosorbent assay or hemolytic assay for detection of complement activity, we demonstrated the ability of the IgM cores to bind C1q and to activate the first step of the classical complement pathway when adsorbed to a surface but with the potentiality to inhibit complement-induced cell lysis when present in solution.

## Results

IgMs have distinct star-shaped structures with a compact Fc-core, comprising the heavy-chain Cμ3, Cμ4 domains, and the terminal tailpieces. Surrounding this core, Fab-Cμ2 arms, composed of the heavy-chain constant domains, Cμ2, Cμ1, the variable VH domain, and paired with the kappa or lambda light-chain constant and variable domains, extend outward with flexible conformations (Fig. S1) (9, 16, 17). We focused on the compact and central assembly of IgMs by restraining the expressed sequence to the last three Cμ2, Cμ3, Cμ4 constant domains and the terminal piece (Fig. S1).

### Protein expression and purification

The expression vector containing the cDNA of the truncated form of IgM μ chain was obtained by mutagenesis and deleting the sequence for heavy variable and Cμ1 domains from a full IgM heavy chain construct previously designed by our team (18). The resulting Fc-core construct was then transfected into the eukaryotic cells HEK293F, either alone to obtain its hexameric oligomeric state (IgM-Fc), or together with an additional vector containing the J chain, to obtain its pentameric form (IgM-Fc-J). Following stable cell line generation, both IgM-Fc constructs were recombinantly expressed and purified from culture media supernatants using a protocol similar to the one established for recombinant full IgMs (19). Samples purified via size-exclusion-chromatography (SEC) were analyzed using semi-native polyacrylamide gel electrophoresis (PAGE) adapted from Vorauer *et al.* (20). At this stage, while IgM-Fc-J samples migrated as a single homogenous band, IgM-Fc samples migrated as a broader band, suggesting the presence of oligomeric heterogeneities (Fig. S2).

Furthermore, the structural integrity and oligomeric states of IgM-Fc samples were also confirmed with negative staining transmission electron microscopy (TEM). As anticipated, both IgM-Fc cores exhibited the characteristic compact, round shapes typical of IgM cores, with recognizable pseudo-symmetrical 6-armed particles for IgM-Fc and asymmetrical 5-armed particles for IgM-Fc-J (Fig. S3).

### Analysis of IgM-Fc core assemblies and their oligomeric distributions by SEC-MALLS, AUC and Mass Photometry

The oligomeric distributions of both samples were further characterized using several biophysical methods after SEC purification.

#### Size-exclusion chromatography coupled to multi-angle-laser-light scattering (SEC-MALLS)

SEC-MALLS detection revealed a homogenous and sharp peak with an experimental mass of 454 kDa for IgM-Fc-J. This value aligns with the theoretical molecular mass of a fully glycosylated pentameric form (399 kDa from amino-acid sequence + 10 to 15% N-glycosylation) (Fig. 1A). Similarly, the IgM-Fc was found to have a single experimental mass of 489 kDa matching the theoretical molecular mass of its hexameric form (460 kDa + 10 to 15% N-glycosylation) (Fig. 1B).

**Figure 1.**
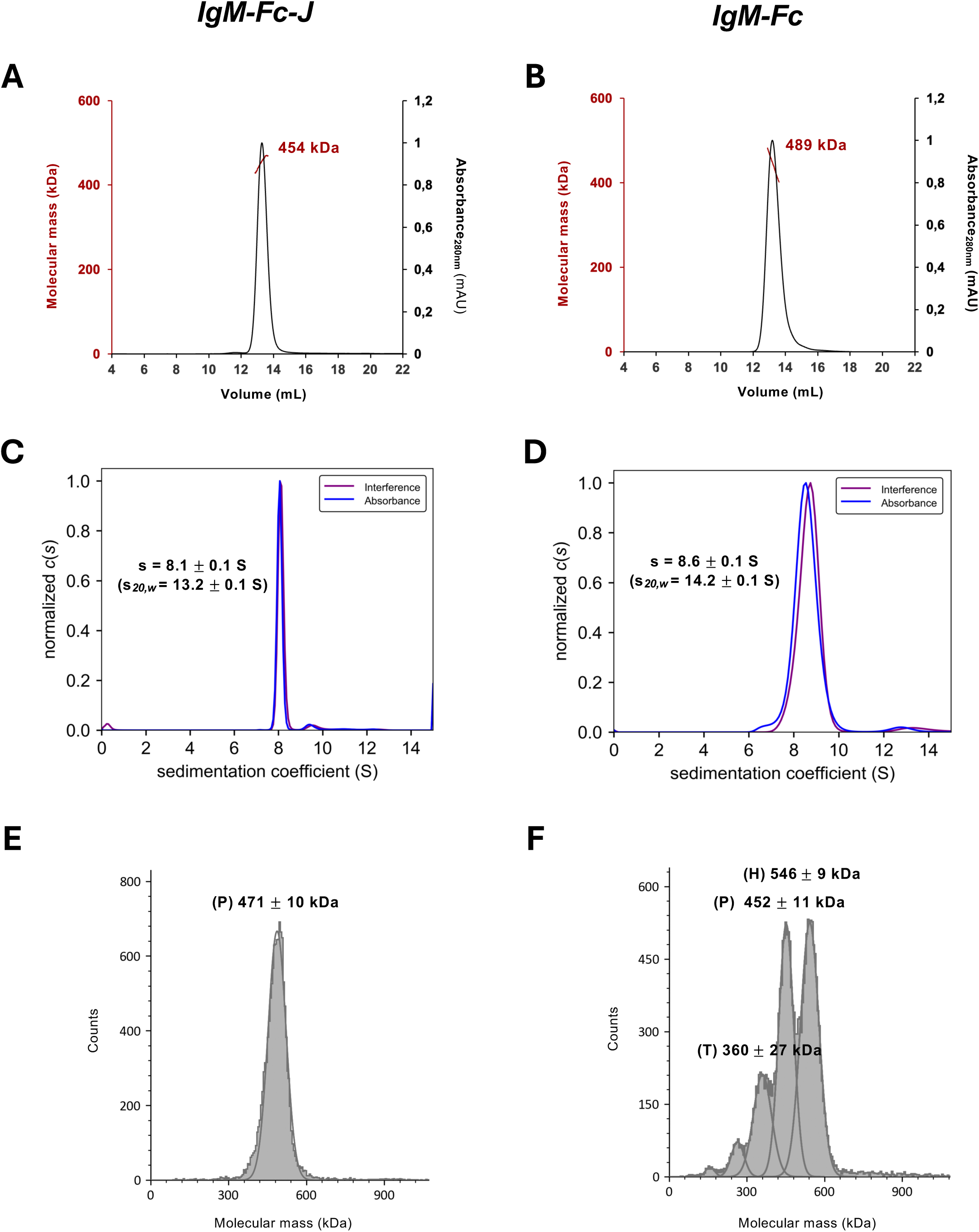
**Analyses of purified IgM-Fc and IgM-Fc-J by size exclusion chromatography coupled to multi-angle light scattering (SEC-MALLS), Analytical ultracentrifugation (AUC) and mass photometry (MP)**. The top panels show the elution profiles of the purified (A) IgM-Fc-J and (B) IgM-Fc monitored by Uv-Vis. absorbance at 280 nm (right ordinate axis) and the molecular mass (left ordinate axis) derived from MALLS, refractometry and UV-Vis measurements. The estimated average molecular masses of the protein peaks are indicated on the graphs. The middle panels show the sedimentation distributions of (C) purified IgM-Fc-J and (D) IgM-Fc. Calculated sedimentation coefficients s and s*_20,w_* in brackets are obtained as described in Materials and Methods and are indicated on the graphs. The bottom panels show the population distributions of purified (E) IgM-Fc-J and (F) IgM-Fc in the SEC peak attributed to the protein fragments. Letters indicate the oligomeric state: H for hexamers, P for pentamers and T for tetramers. Shown analyses are representative of replicate experiments.

#### Sedimentation velocity analytical ultra-centrifugation (sv-AUC)

Since differential oligomeric distributions were previously observed among our full IgM constructs, which have been recombinantly produced using the same procedure (18), and because PAGE heterogeneities were suspected (Fig. S2), further characterization of IgM-Fc and IgM-Fc-J samples was conducted using sedimentation velocity analytical ultra-centrifugation (sv-AUC). Migration in the velocity field of IgM-Fc-J samples revealed a sharp peak at a sedimentation coefficient of about 8.1 S, indicating a highly homogenous pentamer population (Fig. 1C). Similarly, IgM-Fc samples exhibited a single peak at around 8.6 S, albeit broader, indicating a less homogenous hexamer population (Fig. 1D).

#### Mass photometry (MP)

Mass photometry (MP) was further utilized to more precisely assess both IgM-Fc-J and IgM-Fc oligomeric distributions. A single population with an average mass of 471 +/-10 kDa (mean +/- SD over replicates) was observed for IgM-Fc-J (Fig. 1E), aligning with the mass of fully glycosylated pentamers. In contrast, purified IgM-Fc samples displayed a heterogenous population with observed masses of 546 +/- 9 kDa, 452 +-/ 11 kDa and 360 +/- 27 kDa, likely corresponding to hexamers, pentamers without the J-chain, and tetramers, respectively (Fig. 1F).

#### Size-exclusion chromatography coupled to Mass photometry

Due to the limited resolution in SEC separation, a more detailed exploration of the oligomeric distributions of IgM-Fc fragments was conducted by coupling SEC with MP. The approach consisted of combining finely fractionating the size-exclusion peaks with analyzing each fraction using MP. This strategy was applied to IgM constructs lacking the J chain: IgM-Fc from this study and a full recombinant IgM from our previous study, IgM617-HL (18). As anticipated, variations in both oligomeric distributions were observed throughout SEC elution, with a predominant population of hexamers detected in the earliest fractions. This hexameric population decreased along the elution volume in favor of pentamers lacking the J chain, tetramers, and lower oligomer states (Fig. 2A). Interestingly, oligomeric distributions of both IgM-Fc core and full IgM617-HL resemble each other, except for the lower oligomeric population found latter in the elution peak of full IgMs (Fig. 2B).

**Figure 2.**
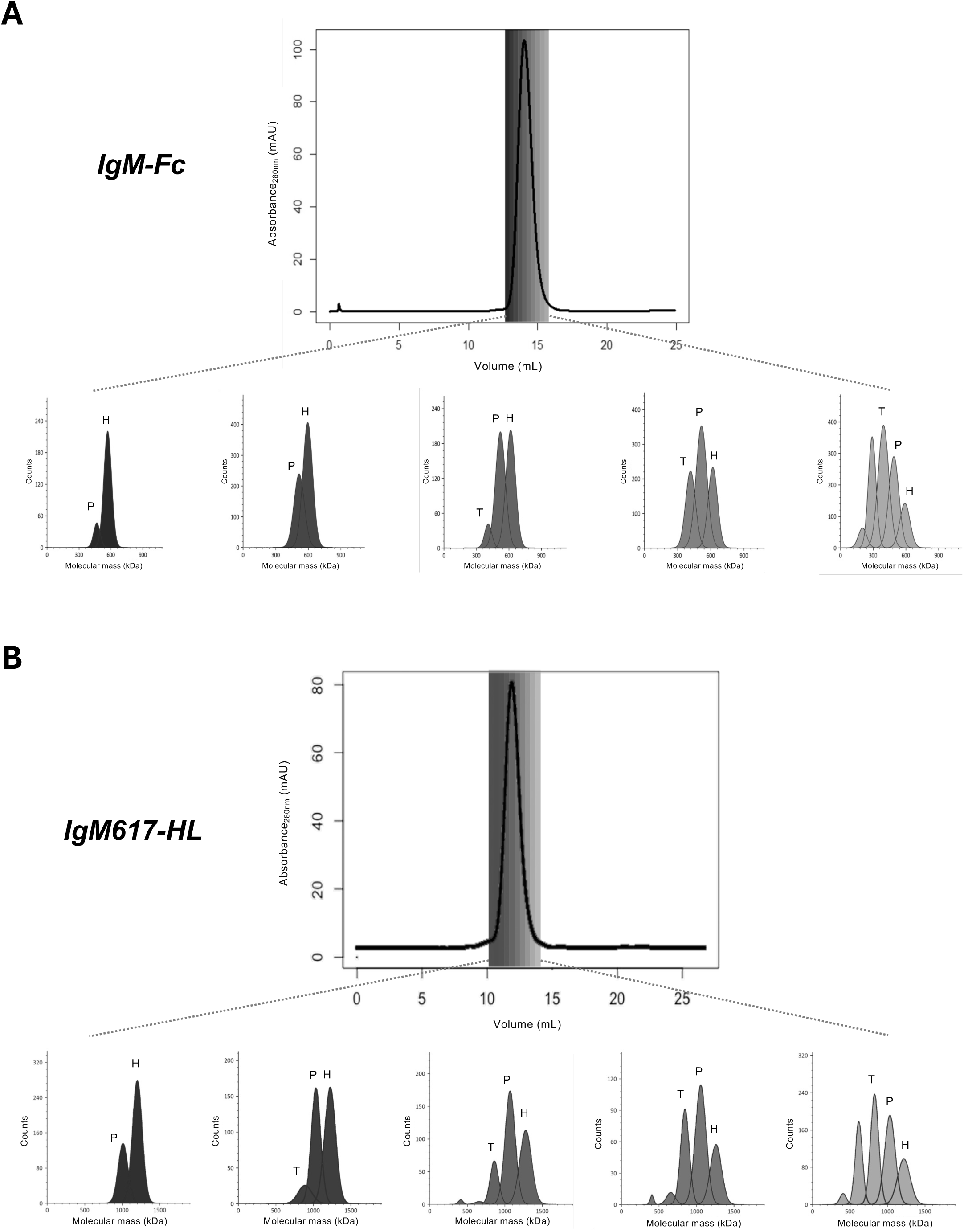
Analysis of purified IgM-Fc (A) and IgM617-HL (B) by size Exclusion Chromatography (SEC) coupled to mass photometry (MP). The upper part of both panels shows the SEC elution profile monitored by Uv-Vis. absorbance at 280 nm of purified IgM-Fc or IgM-Fc-J. The lower part of panels shows population distributions monitored by MP of each individual fraction collected during SEC elutions. Histograms are colored according to the corresponding elution volumes. Letters indicate the oligomeric state: H for hexamers, P for pentamers and T for tetramers.

### In vitro functional analysis of IgM-Fc core assemblies and their oligomeric distributions with ELISA, BLI and hemolytic assays

#### Classical pathway activation by IgM-Fc cores monitored by C4-deposition ELISA

Using our in-house enzyme-linked immunosorbent assay (ELISA) based on the detection of C4b fragment deposition after cleavage of C4 by the C1 complex bound to coated IgM molecules (21), we analyzed the ability of both IgM-Fc and IgM-Fc-J constructs to activate the CP. Notably, both samples triggered C4b deposition in our assays similarly to plasma-derived full IgMs, with a dependency on C1q (Fig. 3A). Furthermore, ELISA using SEC-fractionated IgM-Fc samples, which exhibit different hexamers/pentamers ratios (see above), revealed that the fractions enriched with Fc-core hexamers showed up to five times more C4b deposition compared to fractions with the Fc-core pentamers. A strong correlation with the hexamer and pentamer ratio contained in the samples was also observable with a decrease of C4b deposition along with the decrease of hexamer population in the samples (Fig. 3B). Interestingly, very similar results were obtained with the SEC-fractionated full IgM617-HL (see above), although the effect of the hexamer/pentamer ratio on C4b deposition was less pronounced (Fig. 3C). All together, these results confirm the influence of the polymer distributions of the IgM oligomers on activating the C1 complex and initiating the first step of the CP proteolytic cascade.

**Figure 3.**
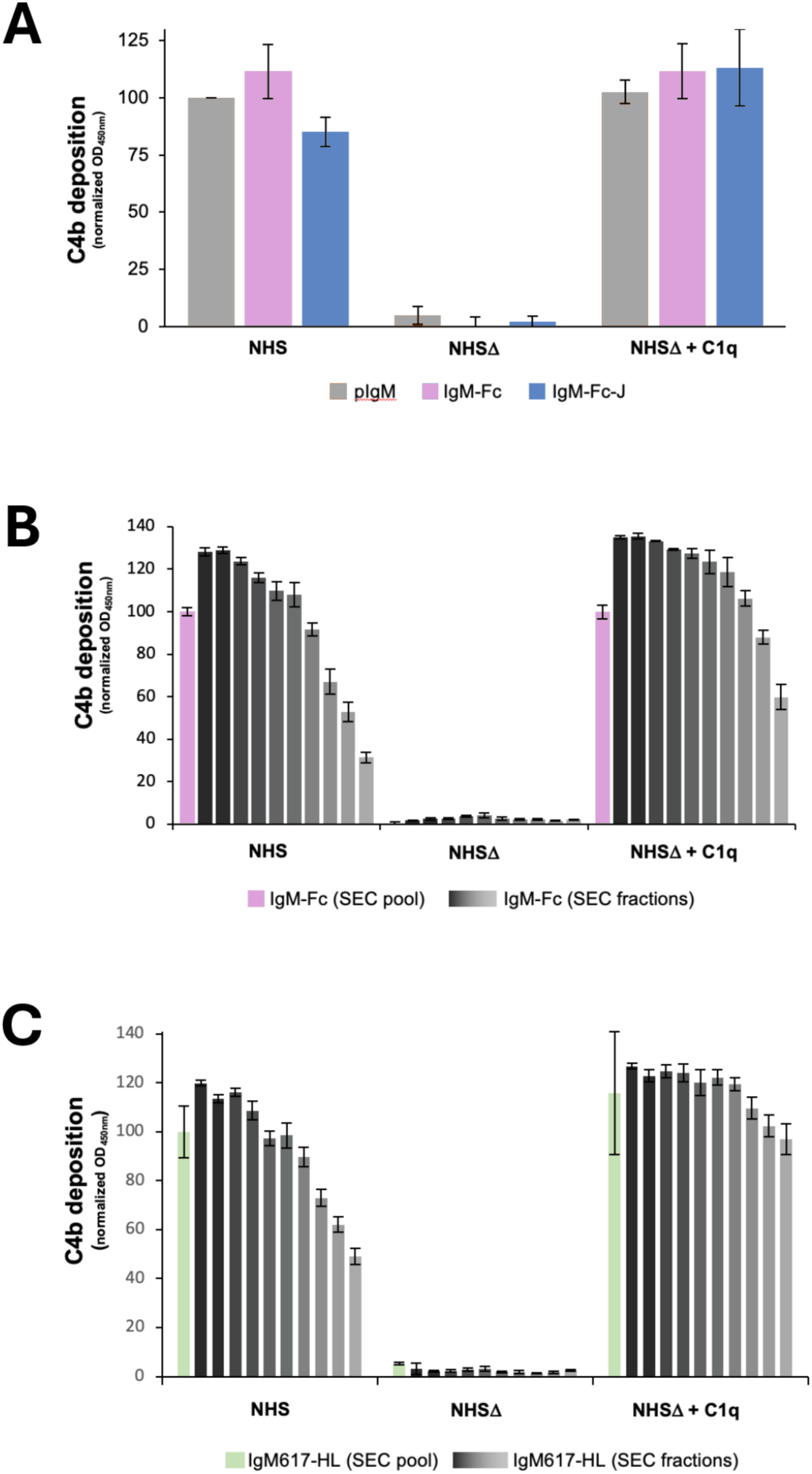
*In vitro* analysis of C1 activation by purified IgM-Fc and IgM-Fc-J. Recombinant IgM-Fc fragments and IgMs purified from plasma were coated on microplate wells and incubated with either Normal Human Serum (NHS), C1q-depleted NHS (NHSΛ), or reconstituted NHS (NHSΛ + C1q). C1 activity was monitored via C4b deposition. Reported values are average of replicated normalized experiments. Errors are obtained as standard deviation between independent replicates. (**A**) C4b deposition comparison between IgMs purified from plasma IgM (pIgM), IgM-Fc and IgM-Fc-J samples obtained in protein SEC peaks (Fig. 1). (**B**) C4b deposition comparison between the IgM-Fc of SEC peak (Fig.1) and the individual SEC fractions (Fig. 2A). (**C**) C4b deposition comparison between the recombinant full IgM617-HL of SEC peak and the individual SEC fractions. (Fig. 2B). Histograms are colored according to the corresponding elution fractions in Fig. 2.

#### Binding of C1q to IgM-Fc cores using BLI

To confirm the dependency of the CP activation on the binding of Fc-core to C1q, biolayer interferometry (BLI) was employed to determine the binding kinetics of plasma-derived C1q to immobilized IgM-Fc and IgM-Fc-J constructs. Since both recombinant Fc-core forms lack an IgM light chain, the BLI strategy using protein L as a capture molecule, which has been applied in our previous study to measure kinetics rate constants of C1q/IgM complex formation, could not be used (18). Therefore, the streptavidin/biotin system was used and optimized to capture biotinylated IgM-Fc and IgM-Fc-J samples (see experimental procedure). Consistent with the observed kinetics of full recombinant IgMs, C1q binding to the IgM cores exhibited a complicated kinetics behavior, which was fitted with a 2:1-heterogenous model to account for the interaction complexity. Under these conditions, the affinities of C1q for both Fc-core forms could be determined in the ten-nanomolar range with a kinetic behavior similar to that observed for the full IgM behaviors previously observed with BLI (Fig. 4 and Table 1) (18).

**Figure 4.**
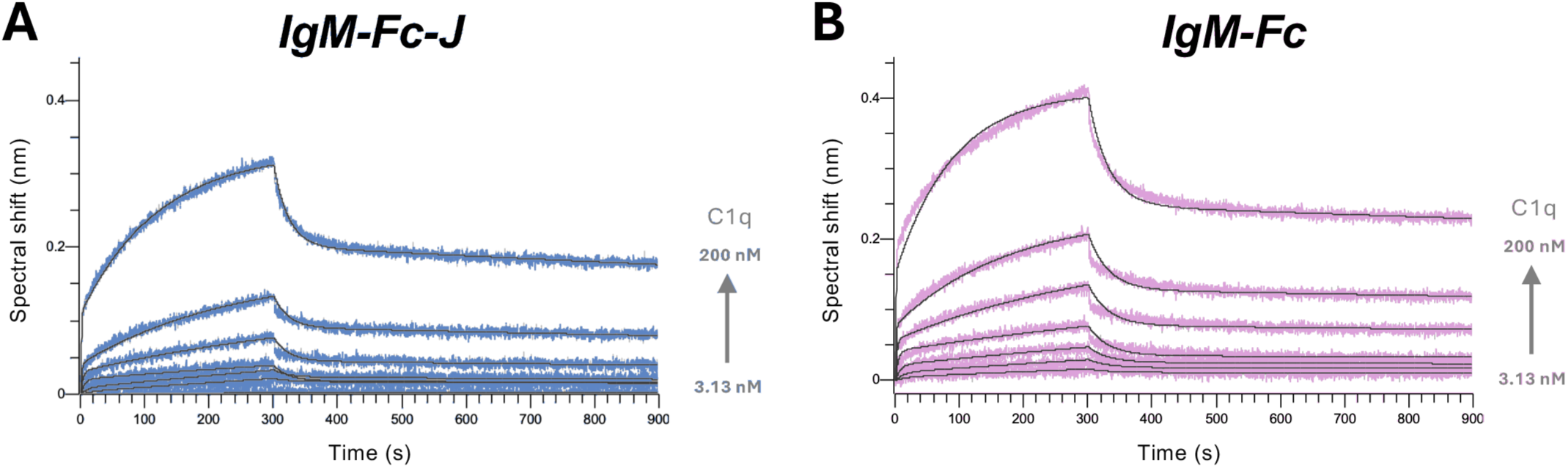
Kinetics analysis of the interaction between C1q and the purified IgM-Fc and IgM-Fc-J. Fc-core fragments were captured on Streptavidin biosensors after pooling SEC fractions containing protein samples. The functionalized biosensors were dipped in wells containing plasma-purified C1q at different concentrations (3.13, 6.25, 12.5, 25, 50, 100 and 200 nM). The binding signals (grey-scaled sensorgrams) were obtained by subtracting the signals from empty biosensors and from zero-concentration samples. Fitted curved are depicted as black lines and kinetics values were obtained by global fitting using a 2:1 heterogenous ligand model and averaging duplicated independent experiments. Shown kinetics analysis are representative of each binding experiment: **(A)** IgM-Fc-J and **(B)** IgM-Fc. Experiments were performed in replicates (n=2).

**Table 1.** Kinetics and affinity parameters of C1 binding to IgM-Fc and IgM-Fc-J obtained by BLI. Experiments were performed in replicates (n=2).

| **Construct** | $k_{a1}$<br>$10^4/\text{Ms}$ | $k_{d1}$<br>$10^{-4}/\text{s}$ | $K_{D1}$<br>$10^{-9} \text{ M}$ | $k_{a2}$<br>$10^6/\text{Ms}$ | $k_{d2}$<br>$10^{-2}/\text{s}$ | $K_{D2}$<br>$10^{-9} \text{ M}$ |
| --- | --- | --- | --- | --- | --- | --- |
| **IgM-Fc** | $2.78 \pm 0.58$ | $3.36 \pm 2.15$ | **$12.1 \pm 0.3$** | $3.33 \pm 0.99$ | $6.13 \pm 5.41$ | **$18.4 \pm 1.8$** |
| **IgM-Fc-J** | $2.06 \pm 0.39$ | $3.87 \pm 1.62$ | **$18.8 \pm 0.1$** | $3.24 \pm 1.64$ | $4.72 \pm 0.66$ | **$14.6 \pm 0.9$** |

#### Complement-induced hemolysis by IgM-Fc core

As both previous assays can be considered as antigen/Fab independent methods to evaluate the functionalities of IgM constructs, we also employed a hemolytic assay to assess the possibility of IgM-Fc forms, as well full IgM617-HLJ, to activate or to inhibit the classical pathway activation in a cellular context (22, 23). This assay measures the lysis of sheep red blood cells (sRBCs) either sensitized or not with a rabbit anti-sRBC antibody (hemolysin), followed by incubation with human plasma as a source of all CP components, supplemented with either IgM-Fc, IgM-Fc-J, or full IgM617-HLJ.

As expected, experiments with full IgM617-HLJ added to plasma showed no effects either on non-sensitized (Fig. S4A) or sensitized cell lysis (Fig. S5A).

When testing the functionality of IgM-Fc and IgM-Fc-J on non-sensitized sRBCs, a surprising and concentration-dependent reduction in hemolysis was observed. Compared to non-sensitized cells incubated with plasma alone, a slight delay of hemolytic activity, as measured by TH50 between 1 to 5 min, was observed upon addition of either IgM-Fc (Fig. S4B) or IgM-Fc-J (Fig. S4C) at a concentration as low as 0.14 nM. Despite no difference was seen in end-point lysis (90 to 99% at 25 min) (Figs S4B and S4C), the addition of IgM-Fc cores led to a diminution of the total complement activity comparable to the residual activity of plasma against non-sensitized cells (Fig. 5). Increasing the concentration of IgM-Fc or IgM-Fc-J up to 4.5 nM resulted in a marked decrease of hemolyzed cells amounts (Figs. S4B & S4C) and a complete loss of total complement activity (Fig. 5).

**Figure 5.**
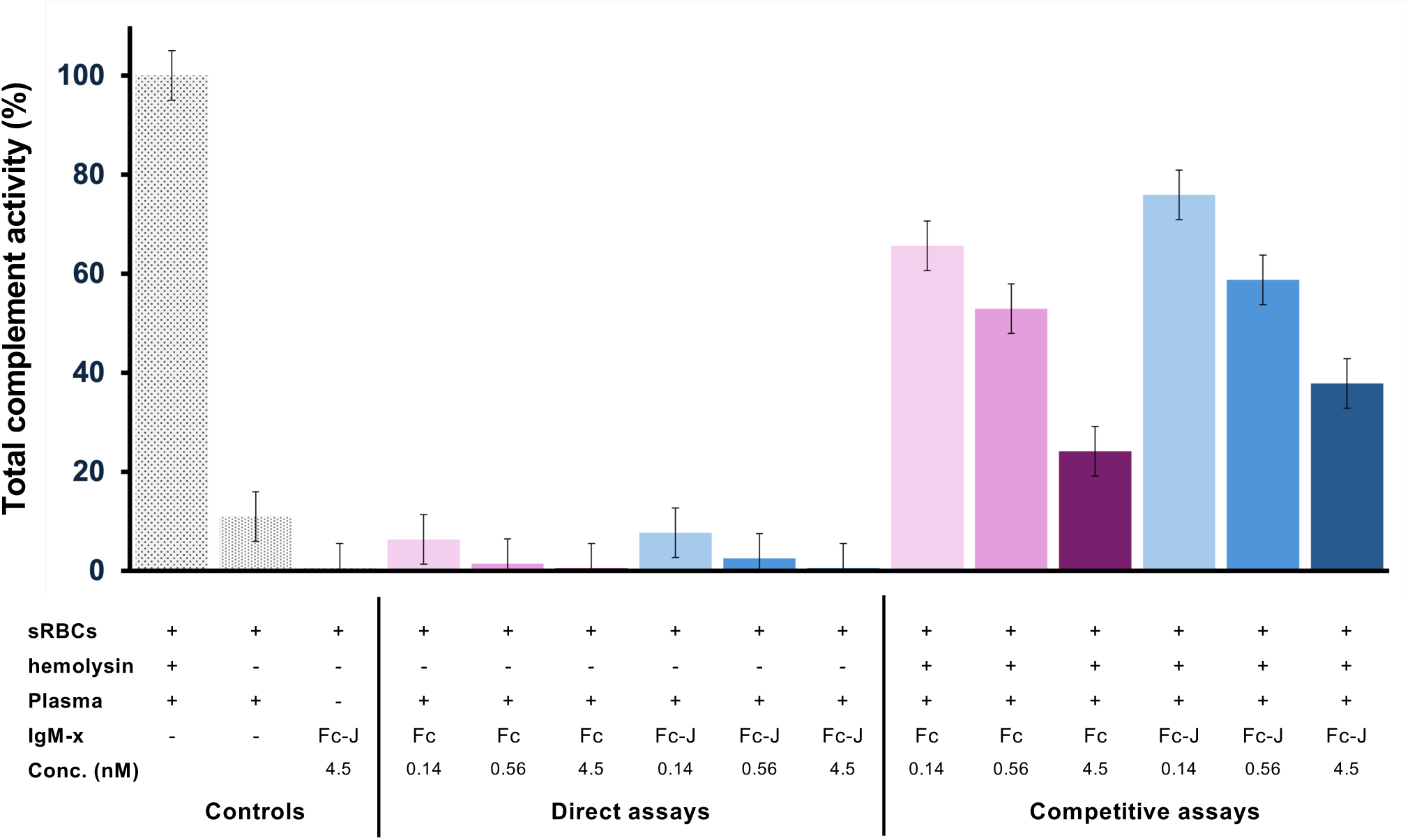
Total complement activities induced by IgM-Fc and IgM-Fc-J. Hemolysis of unsensitized cells (direct assays) or hemolysin-sensitized cells (competitive assays) was measured after mixing either IgM-Fc (pink scale histograms) or IgM-Fc-J (blue scale histograms) at 3 different concentrations with human plasma. Controls (grey scale histograms) were performed the same way but omitting one of the components. Total complement activities are calculated as described in Materials and Methods. Experiments were performed in replicates with 2 different human plasmas.

To investigate the influence of Fc-cores on IgG-induced CDC, a second set of experiments was conducted using hemolysin-sensitized sRBCs and plasma supplemented with varying concentrations of IgM-Fc or IgM-Fc-J. Relevant shifts of TH50 up to 6 min (Figs. S5B & S5C) and decrease of the residual complement activity (Fig. 5) were measurable when adding either one of both constructs up to 4.5 nM. Although decreases were less pronounced than in direct hemolytic assays, the delayed hemolytic rates (5 to 9 min) and residual activities (25 to 38%) tend to be comparable to those measured with non-sensitized cells (9 min and 11%, respectively).

Taken together, the hemolytic assays data suggest that IgM-Fc cores do not enhance CDC *in vitro* as antigen-specific Igs. Interestingly, they might have an inhibitory potential on Igs/complement-induced hemolysis by reducing the total complement activity.

## Discussion

In the pursuit of new biotechnological tools using engineered immunoglobulins, fundamental questions persist particularly regarding the unique attributes of the IgM class. These questions include the ability to produce the original and innovative molecules in recombinant functional forms, to accurately characterize their molecular properties and sample qualities, and to thoroughly understand their mechanisms of action. Such information is indeed crucial for assessing the potential of macro-immunoglobulins in the development of future biological tools or therapeutics. In this study, we focused on expressing and purifying the Fc-core of IgM at the laboratory scale, and on characterizing its quality and functionality using a combination of *in vitro* biochemical and biophysical methods. We expressed two constructs (IgM-Fc-J and IgM-Fc) of IgM μ chain comprising the last three constant domains (Cμ2, Cμ3 and Cμ 4) and the tail piece (tp). By doing so, with or without the J chain in the HEK239F expression system, we expected to obtain either its pentameric or hexameric forms (Fig. S1).

In a previous study, we have observed more heterogeneities in the oligomeric states of full IgMs expressed without the J chain compared to those with the J chain (18). Consequently, we also applied the panel of biophysical analytical methods, including SEC-MALLS, sv-AUC, and MP (Fig. 1), to determine the oligomeric distribution and assess the homogeneity of the two new IgM-Fc constructs. Similar to our findings on full IgMs, IgM-Fc-J core could be purified as highly homogenous pentamers, whereas IgM-Fc still exhibited heterogeneities, notably containing lower assemblies, presumably pentamers lacking the J chains and tetramers in addition to the hexamers. Our results align with most of the previous attempts to produce recombinant full IgM hexamers using different expression systems (24–27). Altogether, these results suggest a crucial role of the J chain in stabilizing the IgM Fc-core, likely in a more favorable manner than a sixth IgM protomer. The structural details by which the additional protomer fails to totally replace the J chain requires further investigation. Nonetheless, our study, along with our previous work on full recombinant IgM models (18, 28), reinforces the importance of extensively studying the polydispersity of large macromolecular complexes like IgMs using advanced biophysical techniques. Conventional methods such as SEC and PAGE, although adapted for IgMs, have not proven completely suitable for detecting heterogeneities in full IgM and IgM-Fc samples. Notably, in the case of IgM-Fc cores, we failed to discern the various oligomers during the purification and quality controls, likely due to the limitations in terms of separation resolution or detection capabilities of those methods. In our studies, MP has emerged as a valuable and reliable method for quickly characterizing the polydispersity of full IgMs and IgM-Fc cores with low sample amounts. To address the limitations of SEC and leverage the advantages of MP, we combined both methods with a strategy involving in depth characterization of the SEC protein elution peak. By applying this approach to both full IgM and IgM-Fc lacking the J chain, we were thus able to isolate different fractions containing different ratios of oligomers, with the first eluted fractions enriched with hexamer populations (Fig. 2). This approach appeared to be faster and easier than previous ones, which have used the combination of sucrose density gradient centrifugation with SDS-PAGE (25, 29).

For assessing complement activation levels, several techniques and assays are available. Hemolytic assays are traditionally used by measuring lysis of sheep red blood cells (sRBCs) opsonized with anti-sheep erythrocyte antibodies (hemolysin) or not sensitized rabbit blood cells. Theses assays are increasingly replaced by liposome immunoassays (LIA), wherein sRBCs are substituted with liposomes coated with antigens and containing a reporter molecule (30). Alternatively, ELISAs using plate coated IgMs, mannose or LPS allow for a more accurate and reproducible quantitation of the different complement components produced after either classical, lectin or alternative pathway activations, respectively (30). During our study, we evaluated the IgM-Fc core ability to activate CP using both ELISA and sRBC hemolytic assays. Notably, our data revealed *in vitro* deposition of C4b in a C1q-dependent manner as detected with ELISA (Fig. 3A). The activation of CDC through CP is thought to be triggered by IgMs only when they are bound to their specific antigens at pathogen surfaces. The antigen binding may be required to induce quaternary structural changes within IgM molecules to allow exposure of C1 binding sites, C1 binding and subsequent proteolytic mechanisms (31). The bias induced by immobilizing/coating IgMs onto *in vitro* surface is well acknowledged for the quantification of complement component depositions and CP activation (32). Our ELISA data with IgM-Fc and IgM-Fc-J constructs confirm this bias. Moreover, they suggest the Fab independency of CP activation in such conditions, which might provoke the necessary structural exposure of the C1q binding motif comprised in the Cμ3 domain and hidden by the Cμ2 domains when IgM are unbound (31, 33).

Furthermore, our method to separate enriched fractions of either IgM-Fc or full IgM hexamers from pentamers and lower oligomeric states, allowed us to explore the differences between the forms in C1 activation. In our previous study, no obvious differences could be observed between pentamer and hexamer constructs. This observation was likely due to the presence of high and variable proportions of pentamers in the purified samples of IgMs lacking the J chain (18). Recently, disparities of up to twofold in *in vitro* C4b depositions have been observed between supposed hexameric and pentameric IgM preparation (34). Our ELISA data clearly demonstrated that the enrichment in hexamers in both IgM-Fc and full IgM-HL samples is associated with an increase of C4b deposition and thus activation of the first CP step (Figs 3B and 3C). Our latest observations are consistent with previous hemolytic assays, which have demonstrated the higher efficacy of hexameric IgMs in inducing CDC compared to pentameric IgMs (25, 35).

The dependency of the activation of CP on direct C1q binding to IgM cores was further confirmed by our new kinetics data obtained via BLI, which showed affinities of C1q for IgM-Fc constructs in the twenty-nanomolar range (Fig. 4). Although the affinity values for Fc cores fall within a similar range as those measured for full IgMs (18, 21, 36), differences of up to one order of magnitude can be easily observed between them. These differences can be explained by the different methods and strategies employed. For instance, in the case of IgM-Fc and IgM-Fc-J constructs, we had to adapt the biotinylating protocol to achieve the molecule capture on the streptavidin BLI biosensors, whereas capturing via protein L has been proven to be more efficient for full IgMs (18). The Fab independency of C1q binding to IgMs and IgGs in optical surface-based methods such as SPR and BLI remains an open question. Indeed, we or others have measured kinetics and affinities between partial or full immunoglobulins and C1q under diverse conditions and without any pre-coated specific antigens. The influence of Igs coating in inducing and stabilizing favorable Ig conformation for C1q binding has been previously suggested (18). Zhou and collaborators also reported that for coated IgGs, the binding of a specific antigen can influence the C1q binding kinetics. While the C1q binding to immobilized Tratuzumab followed by Her2 binding or immobilized preformed Trastuzumab/Her2 complex could be prevented by the presence of HER2 antigen, the opposite was observed for the Adalimumab/TNFα complex. Their observation is strongly related to the different CDC activities of these IgGs (37).

To verify if our IgM-Fc constructs can modulate CDC, hemolytic assays were conducted using either non-sensitized or IgGs-coated sRBCs. With non-sensitized cells, both IgM-Fc and IgM-Fc-J prevent complement induced hemolysis of sheep sRBCs with a concentration dependency: the lysis levels and rates drastically decreased as the amount of Fc cores increase (Fig. S4). Interestingly, the total residual complement activity appeared to be abolished by the presence of the IgM-Fc cores with levels that are even lower than the residual complement activity of plasma on non-sensitized sRBCs (Fig. 5). The ability of the recombinant IgM-Fc cores to inhibit Ig-induced CDC was confirmed with hemolysin-sensitized cells. Although less marked, decreased of the total complement activity was observed when adding IgM-Fcs to plasma to reach residual activities comparable to the one on non-sensitized cells (Fig. 5). Of note, no relevant differences could be reported between inhibitory effects of IgM-Fc and IgM-Fc-J (Figs S4 & S5). Because only IgM-Fc samples obtained prior fine fractionation and hexamer enrichment could be used for those assays, the mixed content of hexamers and pentamers might mask a putative differential effect between oligomeric states.

This capability to modulate and counterbalance the Ig-induced CP activation has been previously documented for IgG-based molecules inspired by the oligomeric structuration of IgMs. For instance, Fc-μTPL309C (15) and GL-2045 (38) have been reported with significant impacts on reducing complement activation in solution, while surface-bound Hexa-Fc (39) retains aptitude to fully activate the complement. Interestingly, some of these Fc constructs designed so far keep the ability to bind C1q, leading to C4b generation, but fail to induce C2 cleavage (15) and/or result in reduced production of C3a and no C5b (15, 38). Thus, it is not surprising to observe CDC inhibition by our IgM-Fc cores in solution, while they still bind C1q and induce C4b deposition when bound to an *in vitro* surface. The detailed molecular mechanism by which the Ig-Fc multimers can block CP remains to be determined. A clue might resid in the absence of the Fab region from all the studied constructs and in our results of assays performed with a non-relevant recombinant full IgM. Contrasting with the IgM-Fc cores, no reduction in hemolytic abilities and in total complement activity could be observed on either non-sensitized or hemolysin-sensitized-cells experiments when full IgMs have been added in plasma (Figs S4A & S5A). As mentioned before, it is well admitted that IgM binding to antigen through Fabs may be required for making C1 complex able to bind to IgM core and activate the complement cascade. One can speculate that the absence of Fab might favor sequestration of the C1 complex in solution by the IgM Fc-cores or Ig-based molecules and thus consumption of the initial complement components such as C4, preventing recruitment of C2 and the subsequent C3 and C5 at the target surface.

In conclusion, we describe the ability of the IgM-Fc core, in both forms, to compete with IgG and to reduce complement-dependent cytotoxicity in hemolytic conditions. This capacity likely relies on the constant fragments of IgM still interacting with C1q as shown by BLI. Surprisingly, as observed with other oligomeric Ig-Fc-based molecules, C1 complex activation and C4b generation are not prevented in ELISA-based activation assays. Taken together, our data question the molecular mechanism of CP regulation in solution. They also reaffirm the promise of new anti-inflammatory candidate molecules that would be inspired by the IgM oligomers to target the complement for the treatment of autoimmune disorders.

## Materials and Methods

### DNA construct generation

Mammalian expression vector for a short construct comprising the Cμ2, Cμ3, Cμ4 domains (Fig. S1) and the terminal tailpiece of the IgM μ chain (pcDNA3.1(+)-Cμ2Cμ3Cμ4Tp) was obtained using site-directed mutagenesis (QuickChange II XL kit, Agilent) to delete the sequence of the variable and Cμ1 domains from the pcDNA3.1(+)-IgM012 H chain vector, a subcloned full length and optimized IgM heavy chain sequence previously published in Chouquet *et al.* (18). The vector construct for the mammalian expression of the human J chain (pcDNA3.1(+)-J-chain) was also described previously in Chouquet *et al.* (18).

### Protein expression

First, to generate a stable HEK293F cell line expressing IgM-Fc, cells were transfected with pcDNA3.1(+)-Cμ2Cμ3Cμ4Tp using 293fectin according to the manufacturer’s protocol (Invitrogen) and generation of the stable line achieved with cultivation in FreeStyle 293 expression medium supplemented with 400 μg/ml G418 (Invitrogen).

These cells were then transfected with pcDNA3.1(+)-J-chain in the same way and the stable transfectants producing IgM-Fc-J were generated using cultivation in medium supplemented with additional 100 μg/ml hygromycin (Sigma-Aldrich). The stable cells expressing each Fc-core construct were then cultivated in Freestyle 293 expression medium under antibiotic pressures and passaged every 3 to 4 days when cell density approached 3.10^6^ cells/ml.

Recombinant expressions of full IgM617-HL and IgM617-HLJ have been previously described in Chouquet *et al.* (18).

### Protein purification

All IgM protein constructs were purified from harvested supernatants according to Hennicke *et al.* (19). Briefly, POROS CaptureSelect^TM^ IgM Affinity Matrix (Thermo Fisher Scientific) was used for affinity chromatography. Culture supernatants were directly applied to packed column and the IgM molecules were eluted with 1 M Arginine, 2 M MgCl_2_, pH 3.5 or pH 4.0. The collected fractions were immediately neutralized with 1 M Tris pH 8.5. The IgM-containing fractions were then pooled, dialyzed against 0.025 M Tris-Base, 0.137 M NaCl, 0.003 M KCl, pH 7.4 and concentrated. For a second purification step consisting of a Size exclusion chromatography (SEC), they were applied on a Superose^TM^ 6 increase 10/300 or 16/600 column (Cytiva) equilibrated in the dialysis buffer at a flow rate of 0.5 mL/min. Proteins were eluted as a single peak. Identification of purified proteins by SDS-PAGE was performed as described in Vorauer-Uhl *et al.* (20) using Native PAGE 3-12% Bis-Tris gels followed by Coomassie Blue staining.

C1q was purified from human serum according to a well-established protocol. Briefly, IgG-ovalbumin insoluble immune aggregates were prepared as described by Arlaud *et al.* (40). After clarification by centrifugation, human serum, obtained from the Etablissement Français du Sang Rhône-Alpes, was incubated with immune aggregates on ice for 45 min. C1/immune complexes were then collected by centrifugation and extensively washed with 20 mM Tris, 120 mM NaCl, 5 mM CaCl_2_ at pH 7.4. Following C1r and C1s release by washing with buffer containing EDTA, C1q was eluted from immune aggregates with 50 mM Tris, 700 mM NaCl at pH 10.0 and recovered by centrifugation. C1q samples were further purified to homogeneity by CM-cellulose chromatography.

### Transmission electron microscopy (TEM)

About 4 μl of diluted IgM-Fc-J or IgM-Fc samples (60 to 80 ng) were applied to a carbon film evaporated onto a mica sheet. The carbon film was then floated off the mica in ∼100 µL 2 % sodium silicotungstate (SST, Agar Scientific) and transferred onto a 400 mesh Cu TEM grid (Delta Microscopies). Images were acquired with a CETA camera on a Tecnai F20 TEM microscope operating at 200 keV.

### Size exclusion chromatography-Multi angle laser light scattering (SEC-MALLS) analyses

SEC combined with online detection by MALLS, refractometry and UV-Vis was used to measure the absolute molecular mass in solution. The SEC runs were performed using a Superose^TM^ 6 increase 10/300 column (Cytiva) equilibrated in 0.025 M Tris-Base, 0.137 M NaCl, 0.003 M KCl, pH 7.4. Fifty μl of protein sample, concentrated to about 1 mg/ml, were injected with a constant flow rate of 0.5 ml/min at room temperature. Online MALLS and differential refractive index detection were performed using a DAWN-HELEOS II detector (Wyatt TechnologyCorp.) with a laser emitting at 690 nm and an Optilab T-rEX detector (Wyatt Technology Corp.), respectively. Averaged molar masses determination was done with the ASTRA6 software, using the “protein conjugate” module. The following refractive index increments and UV-Vis absorbance values were used: *dn/dc* protein = 0.185 mL/g; *dn/dc* glycosylation = 0.15 mL/g; A_280_ (0.1%, 1 cm) = 1.38.

### Analytical ultracentrifugation

Sedimentation velocity analytical ultracentrifugation (sv-AUC) experiments were conducted in an XLI analytical ultracentrifuge (Beckman, Palo Alto, CA) using an ANTi-60 rotor, double channel Ti center pieces (Nanolytics, Germany) of 12- or 3-mm optical path length equipped with sapphire windows and the reference channel being typically filled with the sample solvent. Acquisitions were done overnight at 4°C and at 20000 rpm (32000 g) using absorbance (280 nm) and interference detection. Data processing and analysis were completed using the program SEDFIT (41) from P. Schuck (NIH, USA), REDATE (42) and GUSSI (43) from C. Brautigam (USA), and using standard equations and protocols described previously (44–46).

### Mass photometry

Coverslips (high precision glass coverslips, 24x50 mm^2^, No. 1.5H; Marienfeld) were cleaned by sequential sonication in Milli-Q H_2_O, 50% isopropanol (HPLC grade)/Milli-Q H_2_O, and Milli-Q H_2_O (5 min each), followed by drying with a clean nitrogen stream. To keep the sample droplet in shape, reusable self-adhesive silicone culture wells (Grace Bio-Labs reusable CultureWell^TM^ gaskets) were cut in 4 to 10 segments. To ensure proper adhesion to the coverslips, the gaskets were dried well by using a clean nitrogen stream. To prepare a sample carrier, gaskets were placed in the center of the cleaned coverslip and fixed tightly by applying light pressure with the back of a pipette tip. Protein landing was recorded using a Refeyn One^MP^ (Refeyn Ltd., Oxford, UK) MP system by forming a droplet of each sample at a final concentration of 8 nM in 0.025 M Tris-Base, 0.137 M NaCl, 0.003 M KCl, pH 7.4. Movies were acquired for 120 s (12000 frames) with Acquire^MP^ (Refeyn Ltd., v2.1.1) software using large camera acquisition settings. Contrast-to-mass (C2M) calibration was performed using a mix of proteins with molecular weight of 66, 146, 500, and 1046 kDa. Data were analyzed using Discover^MP^ (Refeyn LTD, v2.1.1) and analysis parameters were set to T1 = 1.2 for threshold 1. The values for number of binned frames (nf = 8), threshold 2 (T2 = 0.25), and median filter kernel (=15) remained constant. The mean peak contrasts were determined in the software using Gaussian fitting. The mean contrast values were then plotted and fitted to a line. The experimental masses were finally obtained by averaging replicates using independent recombinant IgM preparations (2 to 4) and errors were the standard deviation.

### Biolayer interferometry

Biolayer interferometry (BLI) experiments were performed on an OctetRED96e from Sartorius/FortéBio and were recorded with the manufacturer software (Data Acquisition v11.1). IgM-Fc and IgM-Fc-J samples were buffer exchanged against either 0.01 M Na_2_HPO_4_, 0.0018 M KH_2_PO_4_, 0.137 M NaCl, 0.0027 M KCl at pH 7.4 (Phosphate Buffered Saline, PBS) with Zeba Spin Desalting columns (Thermo Fisher Scientific) and were biotinylated using NHS-PEG4-biotin EZ-link kit (Thermo Fisher Scientific) prior to loading. The recommended manufacturer conditions were adapted to achieve a higher coupling degree with one biotin molecule coupled to each protomer (5 to 6 biotin per Fc molecule).

Analyses were performed in 0.2 ml per well in black 96-well plates (96 well PP, Greiner Bio-one) at 25°C at 1000 rpm agitation and using SA (Streptavidin) Biosensors (Sartorius). These tips were pre-wetted in 0.2 ml PBS, 0.02% Tween-20 for 10 min, followed by equilibration in pre-wetting buffer for 120 s. Biotinylated IgM-Fc and IgM-Fc-J samples were applied at concentrations between 50 and 100 μg/ml and loaded for 600 s until reaching a spectrum shift between 3.5 and 4.5 nm, followed by an additional equilibration step of 120 s or more in analysis buffer composed of TBS complemented with 0.002 M CaCl_2_ and 0.02% Tween-20. Kinetics analyses were performed with association phase of plasma C1q monitored for 300 s and with sample diluted in analysis buffer at concentrations between 0 and 200 nM, followed by a dissociation phase in analysis buffer for 600 s. To assess and monitor unspecific binding of analytes, measurements were performed with biosensors treated with the same protocols but replacing ligand solutions with analysis buffer. All measurements were performed in replicates (between 2 to 4) using independent recombinant IgM-Fc and IgM-Fc-J loadings. Kinetics data were processed with the manufacturer software (Data analysis HT v11.1). Signals from reference biosensor and zero-concentration sample were subtracted from the signals obtained for each functionalized biosensor and each analyte concentration. Resulting specific kinetics signals were then fitted using a global fit method and 2:1 heterogeneous ligand model. Reported kinetics parameter values were obtained by averaging the values obtained with replicated assays and reporting errors as the standard deviation.

### Complement activation detection by C4-deposition ELISA

The activation of the classical complement pathway was monitored by an ELISA based on the detection of C4b deposition according to Bally *et al.* (21), Hennicke *et al.* (28) or Chouquet *et al.* (18). Briefly, 200 ng of IgM-Fc or IgM-Fc-J diluted in PBS were adsorbed on a MaxiSorp 96-well plate (Thermo Fisher Scientific) by incubating overnight at 4°C. After washing, unspecific binding was prevented by saturation with PBS complemented with 2% bovine serum albumin (BSA, Sigma Aldrich) for 1 h at 37°C. Replicate wells were then incubated with either Normal Human Serum (NHS) diluted 25 times in a buffer containing 5 mM Veronal, 150 mM NaCl, 5 mM CaCl_2_, 1.5 mM MgCl_2_ at pH 7.4, C1q-depleted serum (NHSΔ, CompTech) diluted 25 times, or NHSΔ diluted 25 times and reconstituted with purified human C1q (4 µg/mL) for 1 h at 37°C. NHS was obtained from the Etablissement Français du Sang Rhône-Alpes (agreement number 21-001 regarding its use in research). The reaction was stopped by washing with a buffer containing 5 mM Veronal, 150 mM NaCl and 5 mM EDTA at pH 7.4. Deposition of cleaved C4 form was detected with a rabbit anti-human C4 polyclonal antibody (Siemens), an anti-rabbit-HRP antibody conjugate (Sigma Aldrich), addition of TMB (Sigma Aldrich) and a Clariostar plate reader (BMG Labtech). Polyclonal IgM isolated from human serum (Sigma Aldrich) was used as control. Blank wells were prepared and processed similarly to wells coated with IgMs but incubated with buffer instead of NHS samples. Reported values were obtained by normalizing each data set (polyclonal IgM/NHS or IgM-Fc or IgM617-HL defined as 100) after blank subtraction and by averaging data obtained in replicated assays using independent recombinant IgM preparations (between 2 to 4); reported errors were the standard deviation of the replicates.

### Hemolytic assays induced by classical complement pathway

Hemolytic assays to evaluate classical pathway activation were performed similarly as previously described in (47). sRBCs (Elitech) were washed three times with DGVB++ buffer (Glucose 2.5%, Veronal 2.5 mM, CaCl2 2.1 mM, MgCl2 0.5 mM, NaCl 72.5 mM, Gelatin 0.05%, pH 7.4), each wash step followed by cell sedimentation (800 g, 10 min, 4°C). Cell concentration was adjusted to 0.5.10^8^ cells/ml. Buffer (for non-sensitized cells) or hemolysin (anti-erythrocyte antibodies, Sigma-Alrich) (for sensitized cells) were then added with a final dilution of 1:5000 and the mixture incubated at 37°C for 15 min. A mixture of 0.9% (v/v) human plasma, non-sensitized or sensitized sRBCs, and dilutions of IgM-Fc, IgM-Fc-J or IgM617-HLJ were incubated at 37°C. Absorbance at 660 nm was recorded during incubations using a spectrophotometer (Safas 190DES) to measure turbidity as rate of erythrocyte lysis. After blank subtraction, defining time zero as the starting time of hemolysis of sensitized RBCs mixed with human plasma and normalization, % of hemolysis values were reported after 25 min incubation and TH50 as the specific time at 50% of lysis. Total residual complement activities are expressed as the ratio: (TH50 of control)/(TH50 of sample)x100. Values were averaged over 2 replicates and 2 different human plasmas. Errors were reported as standard deviations over these replicates.

## Acknowledgments

This work used the Biophysical, AUC, EM and SPR/BLI platforms of the Grenoble Instruct-ERIC center (ISBG; UAR 3518 CNRS-CEA-UGA-EMBL) within the Grenoble Partnership for Structural Biology (PSB), supported by FRISBI (ANR-10-INBS-0005-02) and GRAL, financed within the University Grenoble Alpes graduate school (Ecoles Universitaires de Recherche) CBH-EUR-GS (ANR-17-EURE-0003). We particularly thank Caroline Mas for her assistance at the Biophysical platform, Christine Ebel and Aline Le Roy at AUC platform. The EM facility is supported by the Auvergne Rhône-Alpes Region, the Fondation pour la Recherche Médicale (FRM), the Fonds FEDER and the GIS-Infrastructures en Biologie Santé et Agronomie (IBiSA). This work was not possible without the initial contribution of Polymun Scientific Immunbiologische Forschung GmbH, who provided original cDNA coding for IgM. We would like to thank Cloé Lecluse from IBS for her expertise and help.

This work was supported by the French National Research Agency (grant C1qEffero ANR-16-CE11-0019) and received funding from GRAL, a program from the Chemistry Biology Health (CBH) Graduate School of University Grenoble Alpes (ANR-17-EURE-0003).

## Author contributions

**Conceptualization:** Jean-Baptiste Reiser, Andrea Pinto and Wai Li Ling

**Funding acquisition:** Nicole Thielens, Jean-Baptiste Reiser

**Team supervisions:** Christine Gaboriaud and Renate Kunert

**Investigation and data:** Isabelle Bally, Anne Chouquet, Andrea Pinto, Véronique Rossi, Renate Kunert and Jean-Baptiste Reiser (for protein expression and purification); Andrea Pinto and Jean-Baptiste (for Mass Photometry, ELISA and BLI measurements), Andrea Pinto and Chantal Dumestre-Perard (for hemolytic assays), Andrea Pinto and Wai Li Ling (for electron microscopy).

**Article writing:** Andrea J. Pinto, Wai Li Ling and Jean-Baptiste Reiser.

## Supporting information

This article contains supporting information

## Data availability statement

Raw or processed data are available upon request. All resources produced for the publication are available under a material transfer agreement from CEA or CNRS.

## Abbreviations

Antibody-dependent cellular cytotoxicity: ADCC;
Antibody-dependent cellular phagocytosis: ADCP;
Antigen-binding fragment(s): Fab(s);
Apoptosis inhibitor of macrophage: AIM;
Biolayer interferometry: BLI;
Classical pathway: CP;
Complement-dependent cytotoxicity: CDC;
Enzyme-linked immunosorbent assay: ELISA;
Fragment crystallizable: Fc;
Follicular dendritic cells: FDC;
Heavy chain: H;
Ig(s): Immunoglobulin(s);
IgA(s): Type-A Immunoglobulin(s);
IgG(s): Type-G Immunoglobulin(s);
IgM(s): Type-M Immunoglobulin(s);
Immune complexes: IC;
Joining chain: J;
Light chain: L;
Liposome immunoassays: LIA;
Mass photometry: MP;
Polymeric immunoglobulin receptor: pIgR;
Secretory component: SC;
(Sedimentation velocity) Analytical ultracentrifugation: (sv-)AUC;
Sheep red blood cell(s): sRBC(s);
Size exclusion chromatography: SEC;
Size exclusion chromatography coupled to Multi-Angle Laser Light scattering: SEC-MALLS;
(Sodium dodecyl-sulfate) polyacrylamide gel electrophoresis: (SDS-)PAGE ;
Surface plasmon resonance: SPR;
Tail piece: tp;
Transmission electron microscopy: TEM;

## References

1. Gong, S., and Ruprecht, R. M. (2020) Immunoglobulin M: An Ancient Antiviral Weapon – Rediscovered. Front. Immunol. 11, 1943

2. Jones, K., Savulescu, A. F., Brombacher, F., and Hadebe, S. (2020) Immunoglobulin M in Health and Diseases: How Far Have We Come and What Next? Front. Immunol. 11, 595535

3. Keyt, B. A., Baliga, R., Sinclair, A. M., Carroll, S. F., and Peterson, M. S. (2020) Structure, Function, and Therapeutic Use of IgM Antibodies. Antibodies. 9, 53

4. Matsumoto, M. L. (2022) Molecular Mechanisms of Multimeric Assembly of IgM and IgA. Annu. Rev. Immunol. 40, null

5. Pan, S., Manabe, N., and Yamaguchi, Y. (2021) 3D Structures of IgA, IgM, and Components. Int. J. Mol. Sci. 22, 12776

6. Michaud, E., Mastrandrea, C., Rochereau, N., and Paul, S. (2020) Human Secretory IgM: An Elusive Player in Mucosal Immunity. Trends Immunol. 41, 141–156

7. Shibuya, A., Sakamoto, N., Shimizu, Y., Shibuya, K., Osawa, M., Hiroyama, T., Eyre, H. J., Sutherland, G. R., Endo, Y., Fujita, T., Miyabayashi, T., Sakano, S., Tsuji, T., Nakayama, E., Phillips, J. H., Lanier, L. L., and Nakauchi, H. (2000) Fcα/μ receptor mediates endocytosis of IgM-coated microbes. Nat. Immunol. 1, 441–446

8. Honda, S., Kurita, N., Miyamoto, A., Cho, Y., Usui, K., Takeshita, K., Takahashi, S., Yasui, T., Kikutani, H., Kinoshita, T., Fujita, T., Tahara-Hanaoka, S., Shibuya, K., and Shibuya, A. (2009) Enhanced humoral immune responses against T-independent antigens in Fcα/μR-deficient mice. Proc. Natl. Acad. Sci. 106, 11230–11235

9. Hiramoto, E., Tsutsumi, A., Suzuki, R., Matsuoka, S., Arai, S., Kikkawa, M., and Miyazaki, T. (2018) The IgM pentamer is an asymmetric pentagon with an open groove that binds the AIM protein. Sci. Adv. 4, eaau1199

10. Oskam, N., den Boer, M. A., Lukassen, M. V., Ooijevaar-de Heer, P., Veth, T. S., van Mierlo, G., Lai, S.-H., Derksen, N. I. L., Yin, V., Streutker, M., Franc, V., Šiborová, M., Damen, M. J. A., Kos, D., Barendregt, A., Bondt, A., van Goudoever, J. B., de Haas, C. J. C., Aerts, P. C., Muts, R. M., Rooijakkers, S. H. M., Vidarsson, G., Rispens, T., and Heck, A. J. R. (2023) CD5L is a canonical component of circulatory IgM. Proc. Natl. Acad. Sci. 120, e2311265120

11. Kubagawa, H., Mahmoudi Aliabadi, P., Al-Qaisi, K., Jani, P. K., Honjo, K., Izui, S., Radbruch, A., and Melchers, F. (2024) Functions of IgM fc receptor (FcµR) related to autoimmunity. Autoimmunity. 57, 2323563

12. Ricklin, D., Hajishengallis, G., Yang, K., and Lambris, J. D. (2010) Complement - a key system for immune surveillance and homeostasis. Nat. Immunol. 11, 785–797

13. 13. Cedzyński, M., Thielens, N. M., Mollnes, T. E., and Vorup-Jensen, T. (2019) Editorial: The Role of Complement in Health and Disease. Front. Immunol. 10, 1869

14. Liu, J., Mao, F., Chen, J., Lu, S., Qi, Y., Sun, Y., Fang, L., Yeung, M. L., Liu, C., Yu, G., Li, G., Liu, X., Yao, Y., Huang, P., Hao, D., Liu, Z., Ding, Y., Liu, H., Yang, F., Chen, P., Sa, R., Sheng, Y., Tian, X., Peng, R., Li, X., Luo, J., Cheng, Y., Zheng, Y., Lin, Y., Song, R., Jin, R., Huang, B., Choe, H., Farzan, M., Yuen, K.-Y., Tan, W., Peng, X., Sui, J., and Li, W. (2023) An IgM-like inhalable ACE2 fusion protein broadly neutralizes SARS-CoV-2 variants. Nat. Commun. 14, 5191

15. Spirig, R., Campbell, I. K., Koernig, S., Chen, C.-G., Lewis, B. J. B., Butcher, R., Muir, I., Taylor, S., Chia, J., Leong, D., Simmonds, J., Scotney, P., Schmidt, P., Fabri, L., Hofmann, A., Jordi, M., Spycher, M. O., Cattepoel, S., Brasseit, J., Panousis, C., Rowe, T., Branch, D. R., Baz Morelli, A., Käsermann, F., and Zuercher, A. W. (2018) rIgG1 Fc Hexamer Inhibits Antibody-Mediated Autoimmune Disease via Effects on Complement and FcγRs. J. Immunol. Baltim. Md 1950. 200, 2542–2553

16. Kumar, N., Arthur, C. P., Ciferri, C., and Matsumoto, M. L. (2021) Structure of the human secretory immunoglobulin M core. Structure. 10.1016/j.str.2021.01.002

17. Chen, Q., Menon, R., Calder, L. J., Tolar, P., and Rosenthal, P. B. (2022) Cryomicroscopy reveals the structural basis for a flexible hinge motion in the immunoglobulin M pentamer. Nat. Commun. 13, 6314

18 Chouquet, A., Pinto, A. J., Hennicke, J., Ling, W. L., Bally, I., Schwaigerlehner, L., Thielens, N. M., Kunert, R., and Reiser, J.-B. (2022) Biophysical Characterization of the Oligomeric States of Recombinant Immunoglobulins Type-M and Their C1q-Binding Kinetics by Biolayer Interferometry. Front. Bioeng. Biotechnol. [online] https://www.frontiersin.org/article/10.3389/fbioe.2022.816275 (Accessed May 24, 2022)

19. Hennicke, J., Lastin, A. M., Reinhart, D., Grünwald-Gruber, C., Altmann, F., and Kunert, R. (2017) Glycan profile of CHO derived IgM purified by highly efficient single step affinity chromatography. Anal. Biochem. 539, 162–166

20. Vorauer-Uhl, K., Wallner, J., Lhota, G., Katinger, H., and Kunert, R. (2010) IgM characterization directly performed in crude culture supernatants by a new simple electrophoretic method. J. Immunol. Methods. 359, 21–27

21. Bally, I., Inforzato, A., Dalonneau, F., Stravalaci, M., Bottazzi, B., Gaboriaud, C., and Thielens, N. M. (2019) Interaction of C1q with pentraxin 3 and IgM revisited: mutational studies with recombinant C1q variants. Front. Immunol. 10, 461

22. Fetterhoff, T. J., and McCarthy, R. C. (1984) A micromodification of the CH50 test for the classical pathway of complement. J. Clin. Lab. Immunol. 14, 205–208

23. Costabile, M. (2010) Measuring the 50% Haemolytic Complement (CH50) Activity of Serum. J. Vis. Exp. JoVE. 10.3791/1923

24. Wiersma, E. J., Collins, C., Fazel, S., and Shulman, M. J. (1998) Structural and Functional Analysis of J Chain-Deficient IgM. J. Immunol. 160, 5979–5989

25. Collins, C., Tsui, F. W. L., and Shulman, M. J. (2002) Differential activation of human and guinea pig complement by pentameric and hexameric IgM. Eur. J. Immunol. 32, 1802– 1810

26. Gilmour, J. E. M., Pittman, S., Nesbitt, R., and Scott, M. L. (2008) Effect of the presence or absence of J chain on expression of recombinant anti-Kell immunoglobulin M. Transfus. Med. 18, 167–174

27. Azuma, Y., Ishikawa, Y., Kawai, S., Tsunenari, T., Tsunoda, H., Igawa, T., Iida, S., Nanami, M., Suzuki, M., Irie, R. F., Tsuchiya, M., and Yamada-Okabe, H. (2007) Recombinant Human Hexamer-Dominant IgM Monoclonal Antibody to Ganglioside GM3 for Treatment of Melanoma. Clin. Cancer Res. 13, 2745–2750

28. Hennicke, J., Schwaigerlehner, L., Grünwald-Gruber, C., Bally, I., Ling, W. L., Thielens, N., Reiser, J.-B., and Kunert, R. (2020) Transient pentameric IgM fulfill biological function—Effect of expression host and transfection on IgM properties. PLOS ONE. 15, e0229992

29. Brewer, J. W., Randall, T. D., Parkhouse, R. M., and Corley, R. B. (1994) Mechanism and subcellular localization of secretory IgM polymer assembly. J. Biol. Chem. 269, 17338– 17348

30 Brandwijk, R. J. M. G. E., Michels, M. A. H. M., van Rossum, M., de Nooijer, A. H., Nilsson, P. H., de Bruin, W. C. C., and Toonen, E. J. M. (2022) Pitfalls in complement analysis: A systematic literature review of assessing complement activation. Front. Immunol. 13, 1007102

31. Sharp, T. H., Boyle, A. L., Diebolder, C. A., Kros, A., Koster, A. J., and Gros, P. (2019) Insights into IgM-mediated complement activation based on in situ structures of IgM-C1-C4b. Proc. Natl. Acad. Sci. 116, 11900–11905

32. Zwirner, J., Felber, E., Reiter, C., Riethmüller, G., and Feucht, H. E. (1989) Deposition of complement activation products on plastic-adsorbed immunoglobulins: A simple ELISA technique for the detection of defined complement deficiencies. J. Immunol. Methods. 124, 121–129

33. Arya, S., Chen, F., Spycher, S., Isenman, D. E., Shulman, M. J., and Painter, R. H. (1994) Mapping of amino acid residues in the C mu 3 domain of mouse IgM important in macromolecular assembly and complement-dependent cytolysis. J. Immunol. 152, 1206–1212

34. John, M. M., Hunjadi, M., Hawlin, V., Reiser, J.-B., and Kunert, R. (2024) Interaction Studies of Hexameric and Pentameric IgMs with Serum-Derived C1q and Recombinant C1q Mimetics. Life. 14, 638

35. Randall, T. D., King, L. B., and Corley, R. B. (1990) The biological effects of IgM hexamer formation. Eur. J. Immunol. 20, 1971–1979

36. Bally, I., Ancelet, S., Reiser, J.-B., Rossi, V., Gaboriaud, C., and Thielens, N. M. (2021) Functional recombinant human complement C1q with different affinity tags. J. Immunol. Methods. 492, 113001

37. Zhou, W., Lin, S., Chen, R., Liu, J., and Li, Y. (2018) Characterization of antibody-C1q interactions by Biolayer Interferometry. Anal. Biochem. 549, 143–148

38. Zhou, H., Olsen, H., So, E., Mérigeon, E., Rybin, D., Owens, J., LaRosa, G., Block, D. S., Strome, S. E., and Zhang, X. (2017) A fully recombinant human IgG1 Fc multimer (GL- 2045) inhibits complement-mediated cytotoxicity and induces iC3b. Blood Adv. 1, 504–515

39. Czajkowsky, D. M., Andersen, J. T., Fuchs, A., Wilson, T. J., Mekhaiel, D., Colonna, M., He, J., Shao, Z., Mitchell, D. A., Wu, G., Dell, A., Haslam, S., Lloyd, K. A., Moore, S. C., Sandlie, I., Blundell, P. A., and Pleass, R. J. (2015) Developing the IVIG biomimetic, Hexa-Fc, for drug and vaccine applications. Sci. Rep. 5, 9526

40. Arlaud, G. J., Sim, R. B., Duplaa, A.-M., and Colomb, M. G. (1979) Differential elution of Clq, Cl̄r and Cl̄s from human CT bound to immune aggregates. use in the rapid purification of Cl̄ sub-components. Mol. Immunol. 16, 445–450

41. Schuck, P. (2000) Size-Distribution Analysis of Macromolecules by Sedimentation Velocity Ultracentrifugation and Lamm Equation Modeling. Biophys. J. 78, 1606–1619

42. Zhao, H., Ghirlando, R., Alfonso, C., Arisaka, F., Attali, I., Bain, D. L., Bakhtina, M. M., Becker, D. F., Bedwell, G. J., Bekdemir, A., Besong, T. M. D., Birck, C., Brautigam, C. A., Brennerman, W., Byron, O., Bzowska, A., Chaires, J. B., Chaton, C. T., Cölfen, H., Connaghan, K. D., Crowley, K. A., Curth, U., Daviter, T., Dean, W. L., Díez, A. I., Ebel, C., Eckert, D. M., Eisele, L. E., Eisenstein, E., England, P., Escalante, C., Fagan, J. A., Fairman, R., Finn, R. M., Fischle, W., de la Torre, J. G., Gor, J., Gustafsson, H., Hall, D., Harding, S. E., Cifre, J. G. H., Herr, A. B., Howell, E. E., Isaac, R. S., Jao, S.-C., Jose, D., Kim, S.-J., Kokona, B., Kornblatt, J. A., Kosek, D., Krayukhina, E., Krzizike, D., Kusznir, E. A., Kwon, H., Larson, A., Laue, T. M., Le Roy, A., Leech, A. P., Lilie, H., Luger, K., Luque-Ortega, J. R., Ma, J., May, C. A., Maynard, E. L., Modrak-Wojcik, A., Mok, Y.-F., Mücke, N., Nagel-Steger, L., Narlikar, G. J., Noda, M., Nourse, A., Obsil, T., Park, C. K., Park, J.-K., Pawelek, P. D., Perdue, E. E., Perkins, S. J., Perugini, M. A., Peterson, C. L., Peverelli, M. G., Piszczek, G., Prag, G., Prevelige, P. E., Raynal, B. D. E., Rezabkova, L., Richter, K., Ringel, A. E., Rosenberg, R., Rowe, A. J., Rufer, A. C., Scott, D. J., Seravalli, J. G., Solovyova, A. S., Song, R., Staunton, D., Stoddard, C., Stott, K., Strauss, H. M., Streicher, W. W., Sumida, J. P., Swygert, S. G., Szczepanowski, R. H., Tessmer, I., Toth, R. T., Tripathy, A., Uchiyama, S., Uebel, S. F. W., Unzai, S., Gruber, A. V., von Hippel, P. H., Wandrey, C., Wang, S.-H., Weitzel, S. E., Wielgus-Kutrowska, B., Wolberger, C., Wolff, M., Wright, E., Wu, Y.-S., Wubben, J. M., and Schuck, P. (2015) A Multilaboratory Comparison of Calibration Accuracy and the Performance of External References in Analytical Ultracentrifugation. PLoS ONE. 10, e0126420

43. Brautigam, C. A. (2015) Chapter Five - Calculations and Publication-Quality Illustrations for Analytical Ultracentrifugation Data. Methods Enzymol. 10.1016/bs.mie.2015.05.001

44. Le Roy, A., Nury, H., Wiseman, B., Sarwan, J., Jault, J.-M., and Ebel, C. (2013) Sedimentation Velocity Analytical Ultracentrifugation in Hydrogenated and Deuterated Solvents for the Characterization of Membrane Proteins. in Membrane Biogenesis: Methods and Protocols (Rapaport, D., and Herrmann, J. M. eds), pp. 219–251, Methods in Molecular Biology, Humana Press, Totowa, NJ, 10.1007/978-1-62703-487-6_15

45 Le Roy, A., Wang, K., Schaack, B., Schuck, P., Breyton, C., and Ebel, C. (2015) Chapter Twelve - AUC and Small-Angle Scattering for Membrane Proteins. in Methods in Enzymology (Cole, J. L. ed), pp. 257–286, Analytical Ultracentrifugation, Academic Press, 562, 257–286

46. Salvay, A. G., Santamaria, M., le Maire, M., and Ebel, C. (2008) Analytical Ultracentrifugation Sedimentation Velocity for the Characterization of Detergent-Solubilized Membrane Proteins Ca++-ATPase and ExbB. J. Biol. Phys. 33, 399

47. Dumestre-Pérard, C., Lamy, B., Aldebert, D., Lemaire-Vieille, C., Grillot, R., Brion, J.-P., Gagnon, J., and Cesbron, J.-Y. (2008) Aspergillus Conidia Activate the Complement by the Mannan-Binding Lectin C2 Bypass Mechanism1. J. Immunol. 181, 7100–7105

