## Supplemental data for "The Fc fragment of IgMs binds C1q to activate the first step of the classical complement pathway, while inhibiting complement-dependent cytotoxicity"

### Figure S1

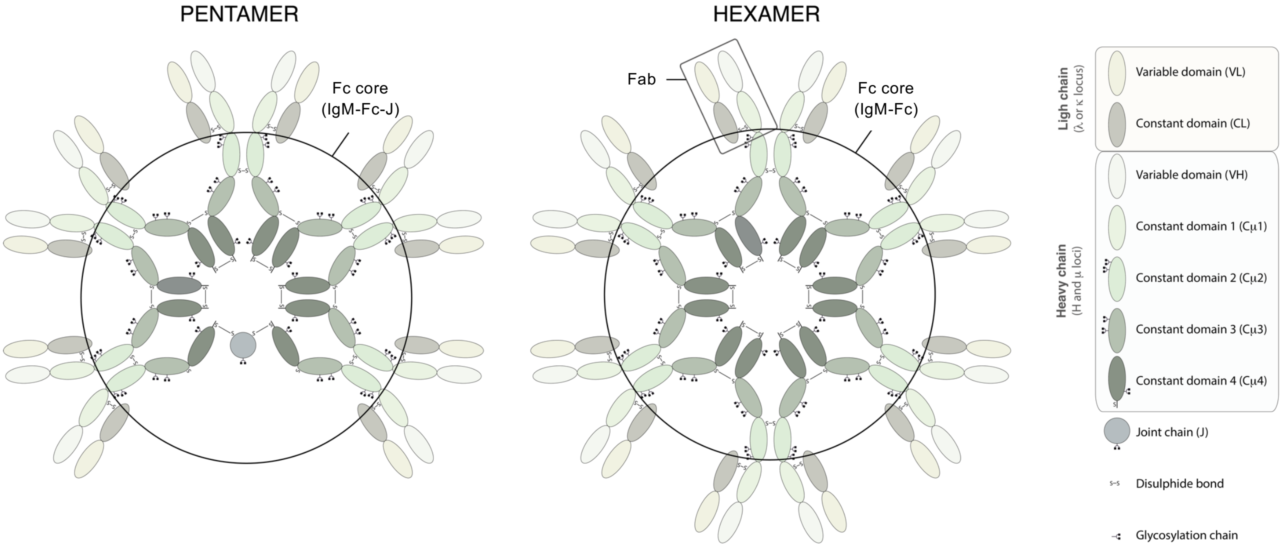


#### Schematic representation of IgMs

### Figure S2

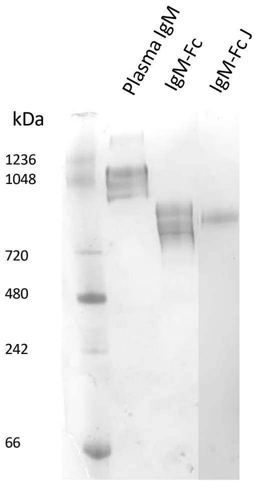


**Semi-native polyacrylamide gel electrophoresis (PAGE) analysis of purified IgM-Fc and IgM-Fc-J samples.** Native PAGE analysis of IgM-Fc and IgM-Fc-J. IgM purified from plasma (Antibodies-onlines GmbH) are shown as control along with native markers (Novex NativeMark$^{TM}$ Unstained Protein Standard).

### Figure S3

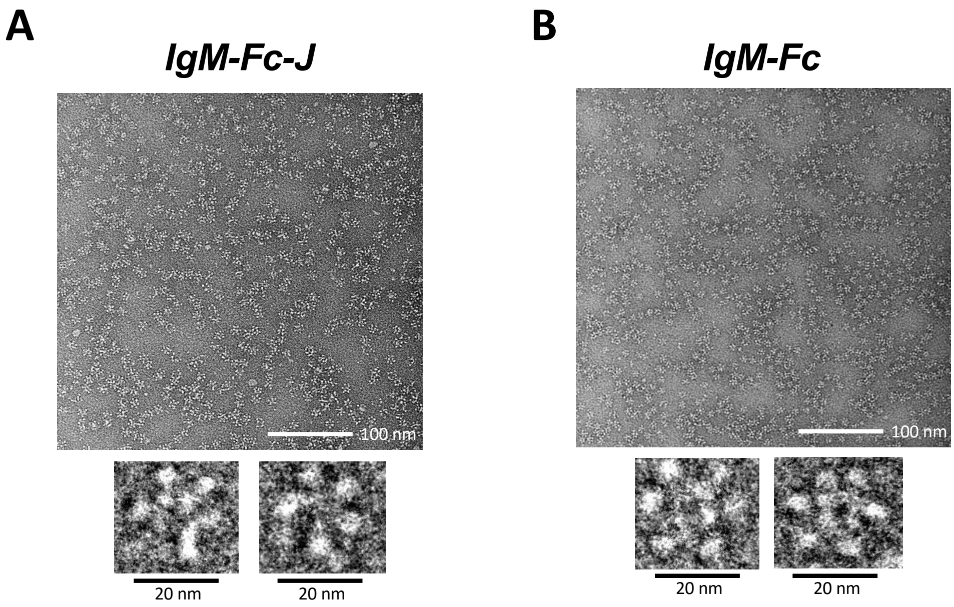


**Non-processed images of recombinant IgM-Fc cores by negative stain transmission electron microscopy.** Representative fields of particles with a 100 nm scale bar are shown on top of each panel with magnified views of some individual molecules shown on the lower part of each panel: **(A)** IgM-Fc-J, **(B)** IgM-Fc.

### Figure S4

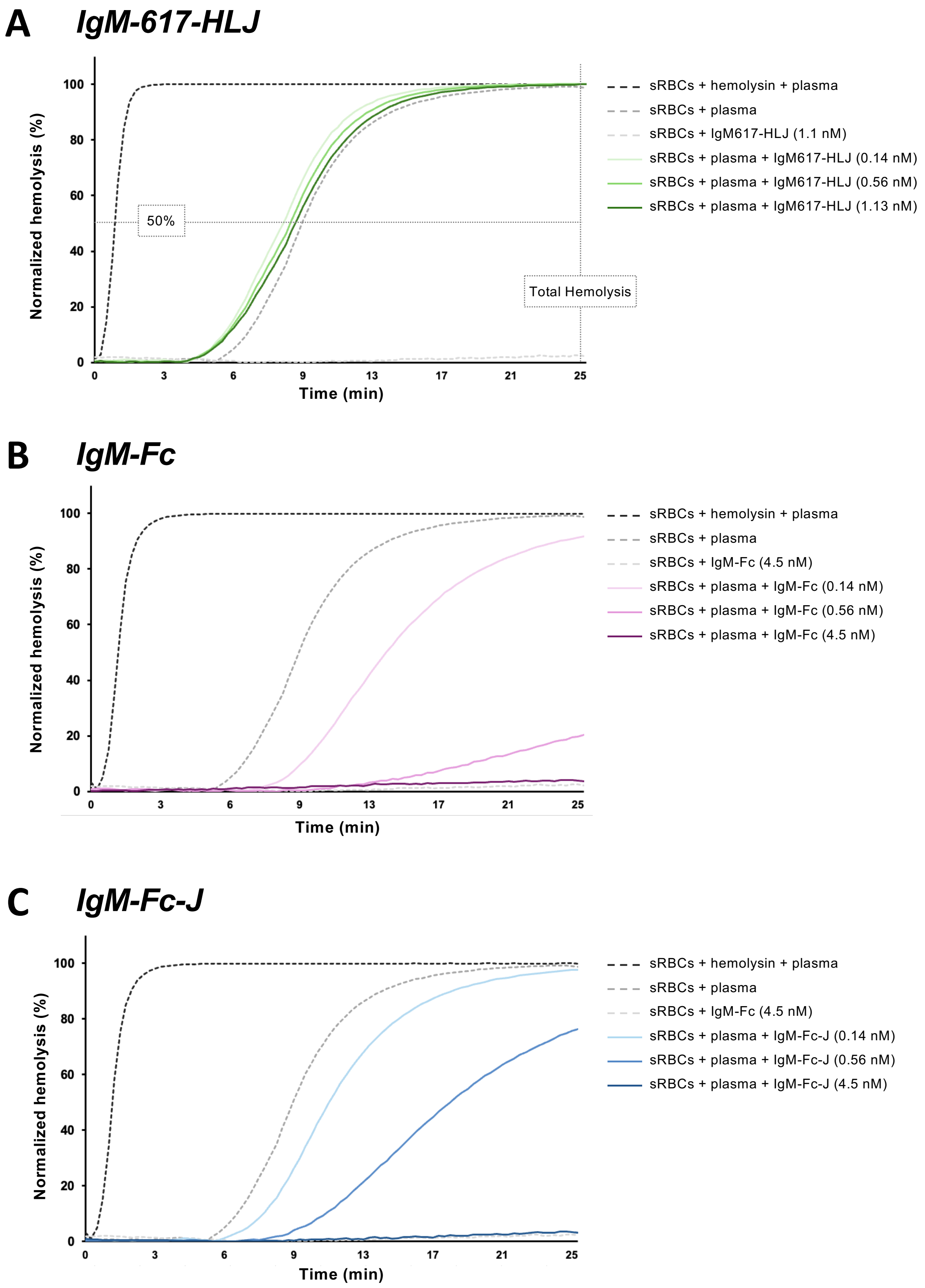


**Time course of direct sRBC hemolysis induced by full IgM and IgM-Fc cores.**

Hemolysis of non-sensitized cells was followed after mixing either (**A**) IgM617-HLJ or (**B**) IgM-Fc or (**C**) IgM-Fc-J with human plasma and incubation of the total mix sRBCs+IgM+plasma for 25 min. Control experiments with sensitized sRBCs with hemolysis+plasma, non-sensitized sRBCs with plasma and non-sensitized sRBS with either IgM617-HLJ or IgM-Fc or IgM-Fc-J at the highest concentration are also reported. Only one representative experiment over replicates is shown.

### Figure S5

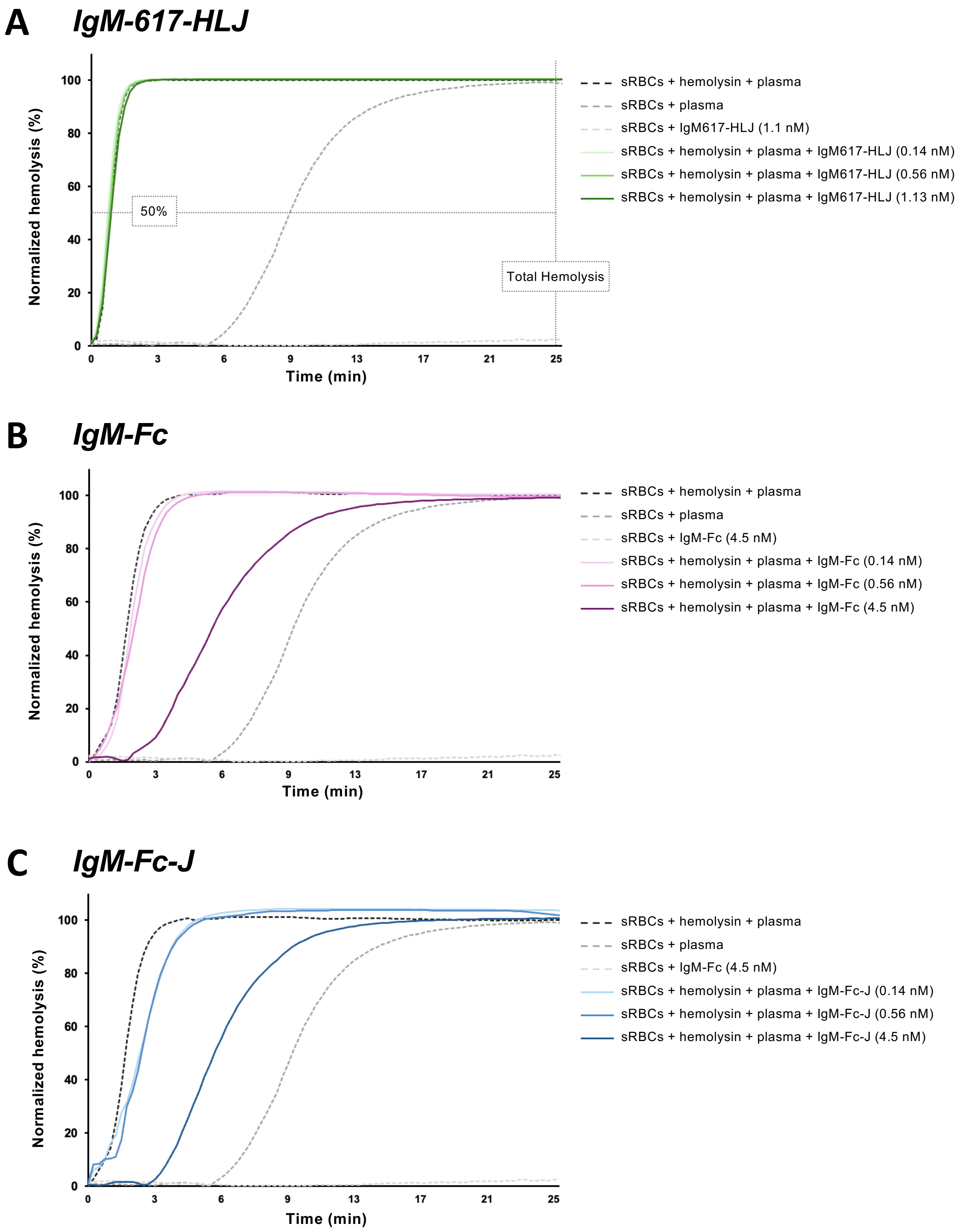


**Time course of competitive sRBC hemolysis induced by full IgM and IgM-Fc cores.**

Hemolysis of hemolysin-sensitized cells was followed after mixing either (**A**) IgM617-HLJ or (**B**) IgM-Fc or (**C**) IgM-Fc-J with human plasma and incubation of the total mix sRBCs+IgM+plasma for 25 min. Control experiments with sensitized sRBCs with hemolysis+plasma, non-sensitized sRBCs with plasma and non-sensitized sRBS with either IgM617-HLJ or IgM-Fc or IgM-Fc-J at the highest concentration are also reported. Only one representative experiment over replicates is shown.
